# Nutritional and Meiotic Induction of Heritable Stress Resistant States in Budding Yeast

**DOI:** 10.1101/390286

**Authors:** Heldder Gutierrez, Bakhtiyar Taghizada, Marc D. Meneghini

## Abstract

Transient exposures to environmental stresses induce altered physiological states in exposed cells that persist after the stresses have been removed. These states, referred to as cellular memory, can even be passed on to daughter cells and may thus be thought of as embodying a form of epigenetic inheritance. We find that meiotically produced spores in the budding yeast *S. cerevisiae* possess a state of heightened stress resistance that, following their germination, persists for numerous mitotic generations. As yeast meiotic development is essentially a starvation response that a/alpha diploid cells engage, we sought to model this phenomenon by subjecting haploid cells to starvation conditions. We find also that haploid cells exposed to glucose withdrawal acquire a state of elevated stress resistance that persists after the reintroduction of these cells to glucose-replete media. Following release from lengthy durations of glucose starvation, we confirm that this physiological state of enhanced stress resistance is propagated in descendants of the exposed cells through two mitotic divisions before fading from the population. In both haploid starved cells and diploid produced meiotic spores we show that their cellular memories are not attributable to trehalose, a widely regarded stress protectant that accumulates in these cell types. Moreover, the heritable stress resistant state induced by glucose starvation in haploid cells is independent of the Msn2/4 transcription factors, which are known to program cellular memory induced by exposure of cells to NaCl. Our findings identify new developmentally and nutritionally induced states of cellular memory that exhibit striking degrees of perdurance and mitotic heritability.

## Introduction

Acquired stress resistance is a phenomenon in which the transient exposure of an organism to a mild environmental stress induces in it the capacity to survive subsequent exposure to what would otherwise be lethal stresses. The occurrence of acquired stress resistance has been documented in numerous species of bacteria, plants, and animals [1–7]. Studies in the budding yeast *Saccharomyces cerevisiae* established the existence of acquired stress resistance in fungi and have contributed greatly to its understanding (for review see [8]). Founding yeast studies showed that following treatment of cells with a moderate level of differing stresses, the exposed cells exhibited resistance to levels of these stresses that were lethal for naïve cells [9]. Interestingly, NaCl exposed cells exhibited cross-stress resistance to exposure to H_2_O_2_. These properties exhibited cellular memory, and persisted in the originally exposed cell after removal of the NaCl inducing stress [10]. Moreover, at some rate, newly born cells inherited this state of acquired stress resistance [10]. Two key features of NaCl acquired stress resistance and cellular memory were demonstrated: 1, topical drug treatments preventing protein synthesis inhibited the acquisition of enhanced stress resistance but had no effect for the cellular memory of this state, and 2, the state of cellular memory was associated with an accelerated transcriptional response to subsequent stress compared with naïve cells, suggesting that the cells also possessed a form of transcriptional memory [9, 10].

Genes exhibiting transcriptional memory in NaCl treated cells include many targets of transcription factors controlling a set of stress resistance genes collectively referred to as the environmental stress response (ESR) [10]. The ESR is a core gene expression program consisting of ~300 upregulated genes and ~600 downregulated genes that is common to cells exposed to diverse environmental stresses and is regulated in part through the paralogous transcription factors *MSN2* and *MSN4* [11]. Indeed, *MSN2*/*4* are required for acquired stress resistance induced by NaCl as well as other stresses [9]. Interestingly, the induction of the ESR by Msn2/4 in naïve cells plays no role in the survival of those cells to the inducing stress [9]. It is thus thought that ESR induction through Msn2/4 serves a preparatory role, acclimating cells to stressful environments through the induction of cellular memory. Although ectopic expression of an Msn2/4 induced catalase gene, *CTT1*, is sufficient to provide resistance to H_2_O_2_, genetic studies suggest that no individual gene accounts for cellular memory [10, 12]. Rather this is a feature that may be brought about collectively by ESR induction, with complex genetic contributions depending on the nature of the inducing and subsequently encountered stresses. Potential unifying physiological mechanisms underlying cellular memory thus remain opaque.

Sporulation represents a developmental occurrence in the yeast lifecycle known to bring about a state of profound stress resistance, a characteristic so far solely attributed to their multilayered spore coat [13]. Here we report that following the shedding of this spore coat during germination and entry into vegetative growth, the resulting cells retain resistance to numerous stresses that are lethal for naïve vegetative cells. Moreover, we show that the resistance of post-germinated cells to hydrogen peroxide (H_2_O_2_) persists in 100% of the population for at least 2 doublings of the culture before fading away, showing that the germinated cells posses cellular memory which is likely inherited by their daughter cells.

We modeled sporulation induced cellular memory by subjecting more experimentally facile haploid cells to glucose starvation, one of the nutritional cues that triggers sporulation, and find that sustained treatments of glucose starvation induce acquired stress resistance and cellular memory comparable to that of spores. Using quantitative single cell assays we show that starvation induced resistance to H_2_O_2_ is inherited by descendants of the originally starved cells through two mitotic divisions. The storage disaccharide trehalose accumulates in sporulated cells as well as in haploid cells that have exhausted the growth media during growth into stationary phase or during glucose starvation [14–16]. As trehalose is a known stress protectant, accumulated trehalose might account for the heritable stress resistant states we describe [8, 17, 18]. We rule out such a role using both biochemical measurements of trehalose levels and genetic studies of trehalose synthase, Tps1. Surprisingly, we also rule out the function of *MSN2/4* and of *CTT1* in the heritable state of elevated H_2_O_2_ resistance in glucose starved cells, thus distinguishing starvation induced cellular memory from previously described forms of cellular memory. Our studies identify new developmentally and nutritionally induced states of cellular memory that exhibit robust perdurance and heritability.

## Results and Discussion

### Yeast spores exist in a stress resistant state that persists in their progeny following germination

Meiotically produced yeast spores exhibit resistance to a profound degree of environmental stress, a trait largely attributable to their spore coat and likely accounting for their remarkable longevity and durability [13]. We sought to determine if spores possess cellular memory contributing to at least some of this stress resistance. To disentangle any cellular memory the spores may possess from the spore coat-conferred stress resistance, we first determined when the spore coat was shed during germination timecourses. Resistance to ether exposure has for long been used as an environmental stress that is specifically attributable to the spore coat and provides a sensitive assay of spore coat integrity [19–21]. To examine germination kinetics, we therefore determined when the loss of spore coat-mediated ether resistance occurred during germination time-courses (Figure 1A). We found that loss of ether resistance was reproducibly observed at the 3-4 hour time point. Therefore, any stress resistance exhibited after these time points cannot be attributable to the spore coat. Of note, the first OD doubling of the germinating culture did not occur until the 6-hour timepoint (Figure 1A), consistent with spore coat shedding preceding the re-entry of the germinated cells into the mitotic cell cycle and vegetative growth.

**Figure 1.**
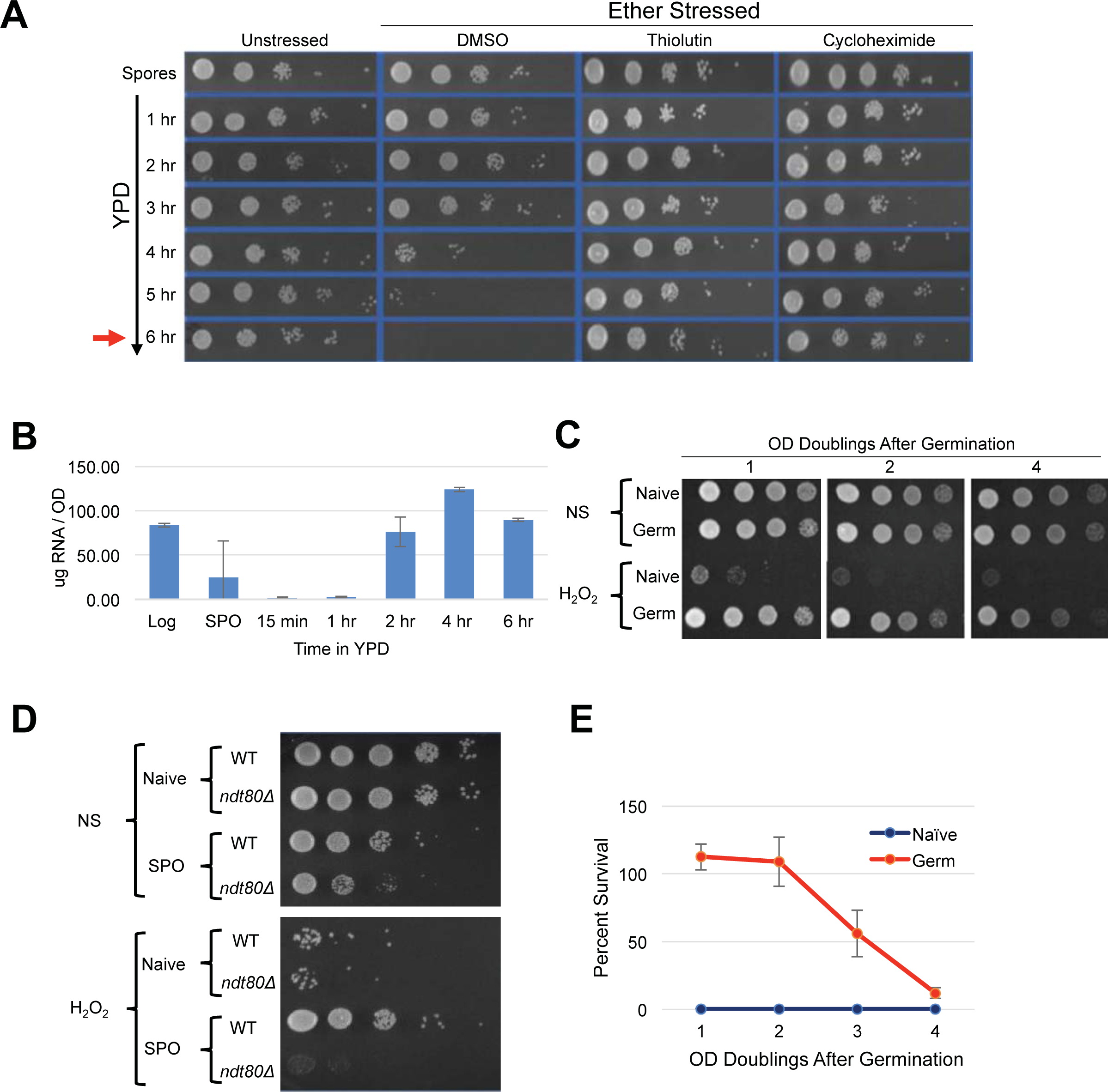
Peroxide resistance persisted following germination. A) Sporulated cells were diluted into YPD media and interrogated with ether treatment. Survival was determined by spotting onto Synthetic complete (SC) plates. The timepoint at which 1 OD doubling was observed is indicated at the 6 hour timepoint and denoted with a red arrow. 10-fold serial dilution of cells is shown. Unstressed - cells germinated without any treatment DMSO – cells treated with control vector for thiolutin and cycloheximide. Thiolutin – cells treated with 10ug/ml of thiolutin. Cycloheximide – cells treated with 2ug/ml cycloheximide. B) Extractable RNA levels during germination of cells cultured in YPD are shown. Samples were taken from mid-log cells (Log), spores (SPO), and germinating cells in YPD at 15 min, 1 hr, 2hr, 4hr, and 6 hours after release from sporulation. C) 10-fold serial dilution of H_2_O_2_stress resistance spot assay; Naive represent cells maintained at mid-log and Germ represent cells growing after germination in YPD media. D) 10-fold serial dilution of H_2_O_2_stress resistance spot assay; WT and *ndt80Δ/∆* cells were grown in YPD (Naive) or sporulation media (SPO) and with (H_2_O_2_) or without (NS) peroxide treatment. E) CFU quantification of resistance to 4 mM H_2_O_2_after germination is plotted. Error bars represent 1 standard error of the mean from 3 biological replicates.

A previous study demonstrated the importance of protein synthesis for germination by exposing germinating cells to the protein synthesis inhibitor, cycloheximide [22]. We confirmed this finding using our ether sensitivity acquisition assay (Figure 1A). To test the hypothesis that new transcription is also required for germination, we reintroduced spores into rich media in the presence of the yeast RNA polymerase II inhibitor thiolutin. Similarly to as we observed with cycloheximide treated spores, thiolutin treatment prevented the acquisition of ether sensitivity (Figure 1A), indicating that new transcription may be required for this defining step of germination. As thiolutin has been reported to also inhibit protein synthesis, we determined when bulk RNA synthesis likely commenced during germination to further test this hypothesis [23]. Consistent with the conclusion that new transcription was required for spore coat shedding, we find that bulk extractable RNA levels dramatically increased at the 2-hour timepoint of germination, at least 1-2 hours prior to spore coat shedding (Figure 1B).

To determine if post-germinated cells retained resistance to other environmental stresses, we exposed germinating cultures that had undergone one full OD doubling to levels of H_2_O_2_, EtOH, or NaCl that were lethal for naïve cells and assayed for survival using spot test assays. We found that these post-germinated cells exhibited striking degrees of resistance to all three of these environmental stresses (Figure 1C and S1A). Because the H_2_O_2_ exposure assay was the most acutely acting, it represented the most facile way to investigate this stress resistant state further. We first determined that an *ndt80∆/∆* strain, which causes meiotic arrest [24], failed to exhibit H_2_O_2_ resistance, confirming that meiotic development was required for cellular memory (Figure 1D). Remarkably, all cells produced following spore germination exhibited H_2_O_2_ resistance, and this characteristic persisted in the post-germinated culture for more than 4 OD doublings before fading from the population (Figure 1C). We quantified the perdurance of this state using plating experiments that permitted precise counts of large numbers of colonies and found that 100% of the cells retained H_2_O_2_ resistance through at least 2 OD doublings before gradually fading from the population (Figure 1E).

These findings establish a new characteristic of yeast gametes: cellular memory. Figure 2A diagrams the developmental dynamics in ether resistance, RNA accumulation, H_2_O_2_ resistance, and trehalose accumulation (discussed below) through sporulation and germination. We next inspected germinating populations microscopically to determine when mating and zygote formation occurred and found that these processes were highly asynchronous (Figure 2B). Thus, although the processes we document during germination itself are quite synchronous with reasonably precise timing, following germination, the cells’ re-entry into the cell cycle is complexly intertwined with mating and zygote formation, complicating further investigation of the heritability of spore cellular memory.

**Figure 2.**
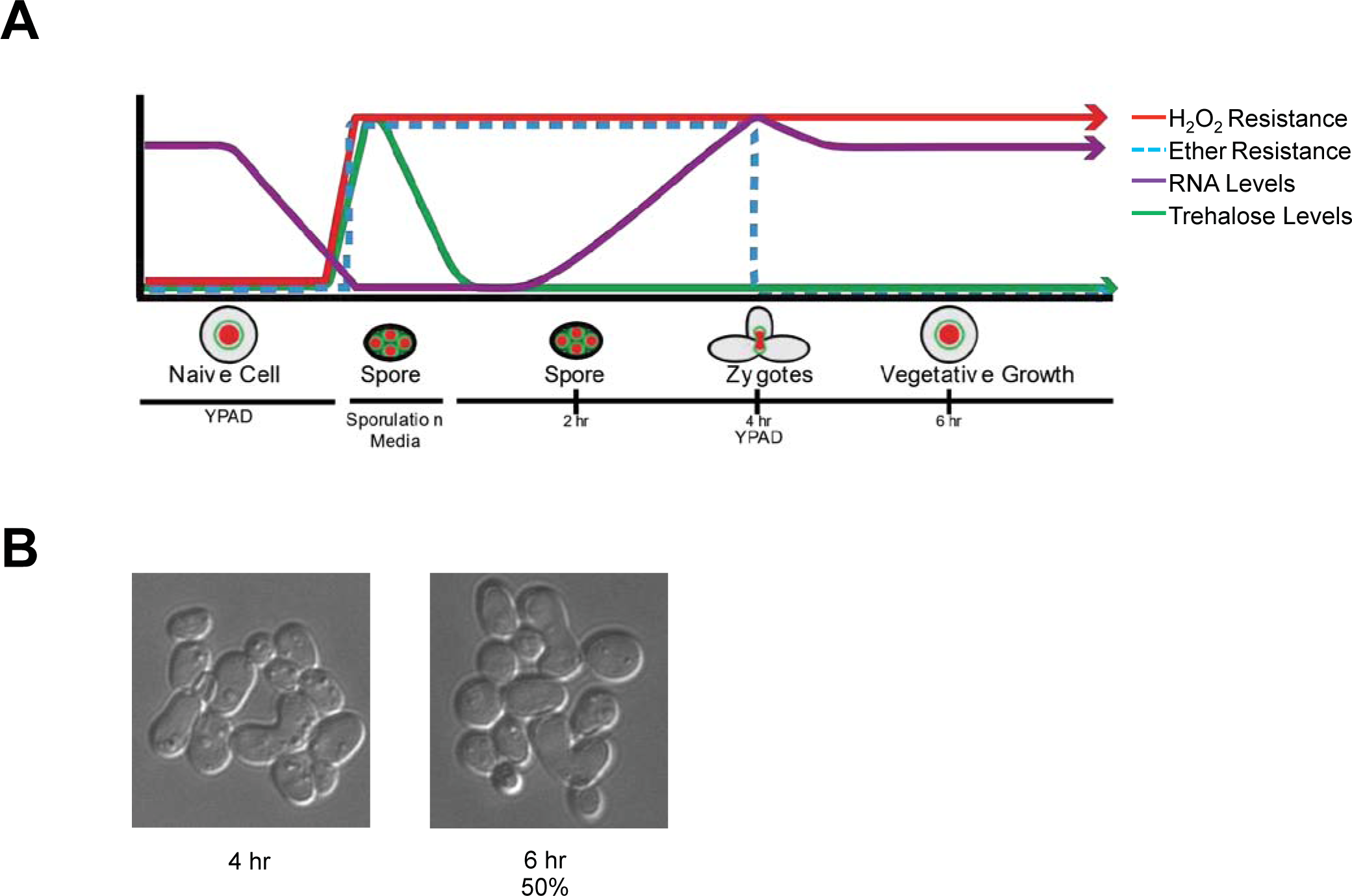
Developmental transitions and cellular memory of germinating cells. A) A diagram depicting the changes in observed H_2_O_2_, ether resistance, trehalose accumulation, and extractable RNA levels through sporulation and germination is shown. B) Representative pictures of germinating cells in YPD media at 4 and 6-hour time points. At the 6-hr time point 1 OD doubling was measured via a spectrometer and ~50% of cells were zygotes as determined by microscopic inspection.

### Glucose starvation of haploid mitotic cells induced a heritable state of cellular memory comparable to that of meiotically produced spores

The conventional protocol for induction of synchronous and efficient sporulation that we use employs an overnight pre-culture in rich media containing acetate (YPA), which differs from the growth conditions of naïve cells (YPD: rich media + dextrose). We found that both WT and *ndt80∆/∆* diploids grown in YPA exhibited robust acquired stress resistance, superficially equivalent to that of germinated spores (Figure 3A). As meiotically arrested *ndt80∆/∆* cells that were incubated in sporulation medium showed no H_2_O_2_ resistance (Figure 1D), our results suggest that the acquired stress resistance of *ndt80∆/∆* cells pre-grown in YPA was somehow lost during their meiotic arrest. Cells arrested in meiosis due to loss of *NDT80* can return to mitotic growth following their dilution into YPD medium [24]. Curiously, meiotically arrested *ndt80∆/∆* cells that had returned to mitotic growth exhibited enhanced H_2_O_2_ resistance (Figure 3B). A parsimonious explanation for these findings is that meiotically arrested cells are simply more vulnerable to environmental stress. In this scenario, the YPA-induced stress resistance of *ndt80∆/∆* cells persists in meiotically arrested cells, but is masked in them by the vulnerable condition of their arrest; upon return of these cells to conditions that relieve meiotic arrest (YPD), their stress resistant state becomes once again detectable (Figure 3B). It seems likely that at least some of the acquired stress resistance and cellular memory of sporulated cells may be induced merely by the nutrient pre-conditions of sporulation (YPA), and are entirely independent of sporulation. Indeed, we determined that MAT**a** haploid cells also exhibited robust acquired stress resistance upon growth in YPA (Figure 3C).

**Figure 3.**
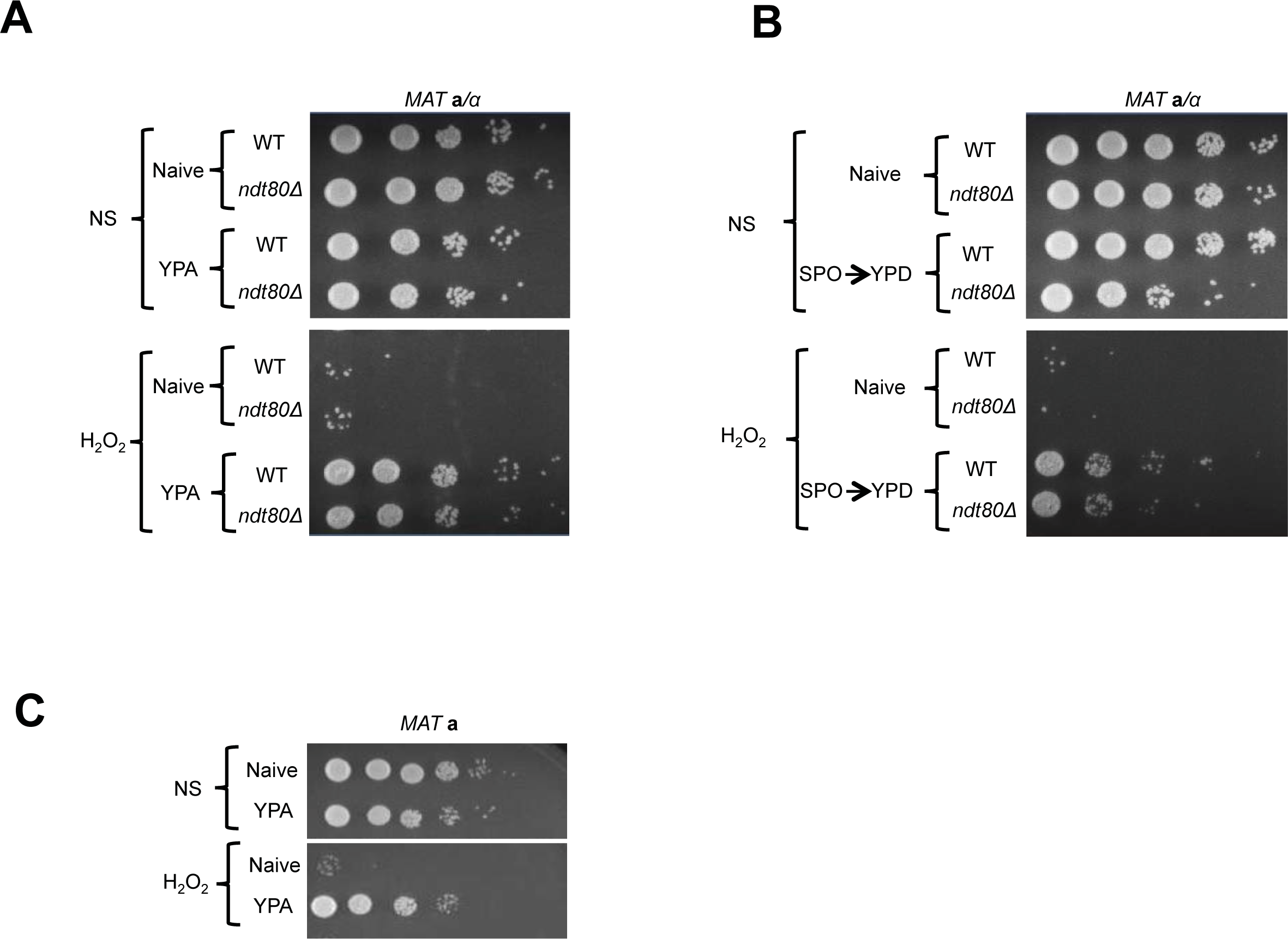
YPA media induced acquired stress resistance. A) WT or *ndt80∆/∆* a/alpha diploid cells were grown in YPA media and subjected to H_2_O_2_ stress. 10-fold serial dilution spot assay of Naïve or YPA grown cells are shown. B) WT and *ndt80∆/∆* strains were incubated in sporulation media to induce spore formation or meiotic arrest respectively. Following dilution of these cells into YPD and incubation for xx hours (SPO → YPD), resistance to H_2_O_2_ was assayed using spotting tests. C) 10-fold serial dilution spot assay of haploid cells grown in YPA media subjected to H_2_O_2_ stress or not is shown.

As mentioned above, YPA media replaces glucose with the non-fermentable carbon acetate. A previous study showed that following 4 days of glucose starvation in synthetic media, yeast cells exhibited resistance to heat and H_2_O_2_ treatments [16]. To determine if shorter intervals of glucose withdrawal in rich YP media was sufficient to induce acquired stress resistance, we interrogated the stress resistance of MAT**a** haploid cells that had been subjected to glucose starvation for varied durations. Following 1 hour of glucose starvation, a slight degree of acquired stress resistance was apparent, though only a small number of cells in the population exhibited this characteristic (Figure 4A). In contrast, 6 and 12-hour regimens of glucose starvation robustly induced acquired stress resistance to H_2_O_2_, EtOH, and NaCl (Figure 4A and S1B). Moreover, we found that the acquired stress resistance in both 6 and 12-hour glucose starved cells persisted for multiple OD doublings after these cells were inoculated back into YPD, confirming that this state exhibited cellular memory (Figure 4A). To determine if starvation induced acquired stress resistance was mitotically heritable, we performed quantitative survival assays and found that this state was indeed inherited by all of the cells through more than 1 OD doubling (Figure 4B). The kinetics of the loss of stress resistance suggested that most, but likely not all, cells inherited this state through a second OD doubling before fading in the population (Figure 4B). These findings thus identify glucose starvation as an inducing condition of acquired stress resistance and cellular memory that is nearly equivalent to that exhibited by spore progeny. Unlike in post-germinated cells, our studies of glucose starved haploid cells permitted an unambiguous assessment of the heritability of this state. We confirm that the glucose starvation induced state of enhanced stress resistance was inherited by mitotic daughter cells through at least one, and in most cases two, divisions.

**Figure 4.**
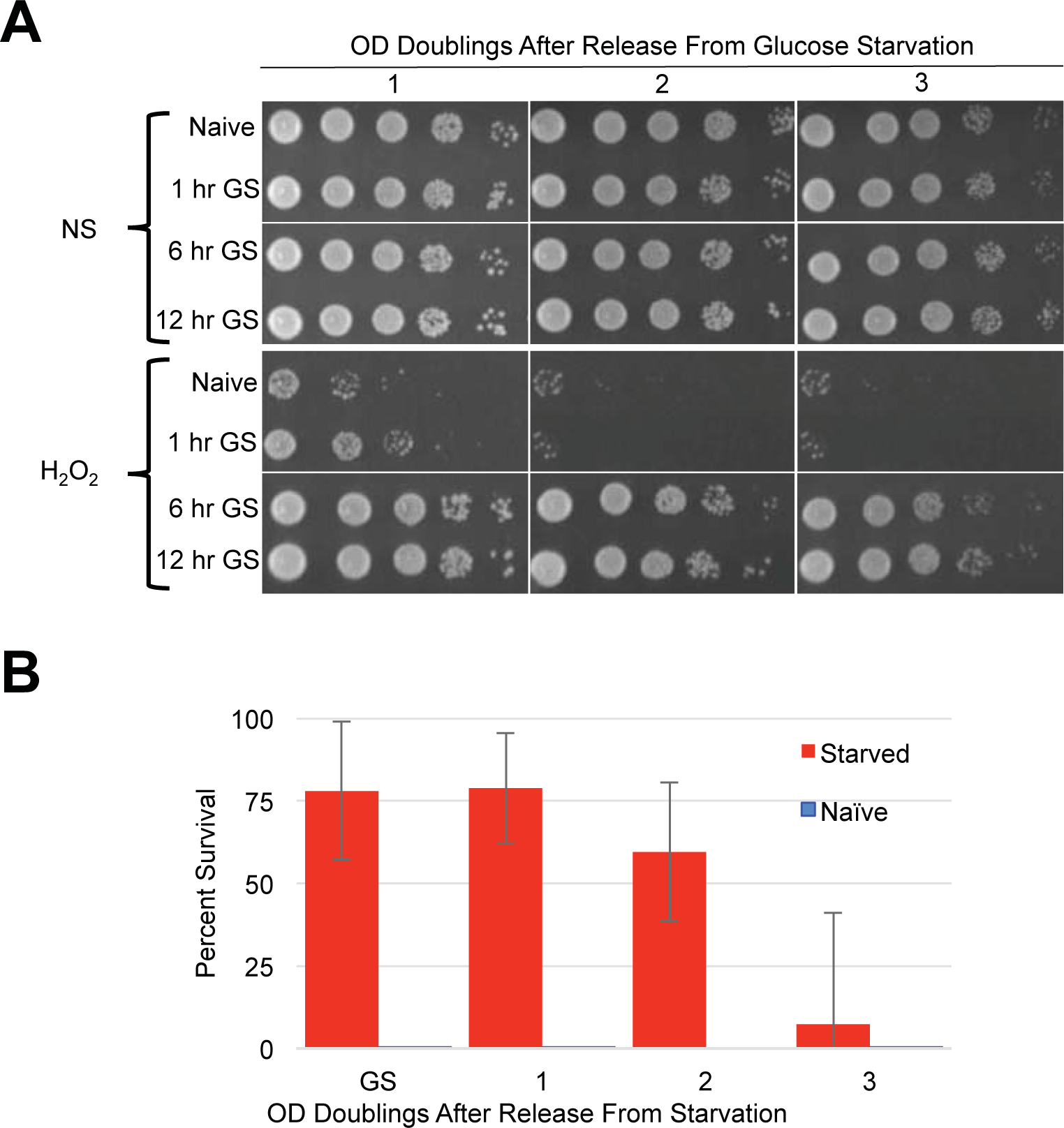
Cellular memory of glucose starved cells. A) Cells grown to log phase in YPD media were harvested and incubated in YP media lacking glucose for varied times. Following glucose withdrawal, cells were subjected to H_2_O_2_ stress and survival was assayed using spotting assays. B) Survival of cells released from glucose starvation and exposed to H_2_O_2_ was quantified. Error bars represent 1 standard error of the mean from 3 biological replicates.

### Starvation induced cellular memory does not require trehalose accumulation or *TPS1* function

A simple potential mechanism underlying the starvation induced stress resistant states and their apparent heritability could be the accumulation of a cyto-protectant that can be distributed through mitotic divisions. The primary candidate for such a cyto-protectant is the disaccharide trehalose, which is known to accumulate in spores and cells that have reached stationary phase following media exhaustion [14–16]. Indeed, trehalose has been associated with cellular protection from a wide variety of stress such as desiccation, heat, and oxidative agents [8, 17]. In dividing stress-sensitive cells, steady state levels of trehalose are negligible, though recent findings suggest that cycles of trehalose production and break down prevent glycolytic catastrophe in them [25–28]. Other recent findings have called into question the stress protective roles of trehalose *per se*, and suggest some unknown function of the trehalose synthase enzyme, Tps1, in stress protective mechanisms [29, 30].

Similarly to what has been described previously, we detected dramatic accumulations of trehalose in spores, stationary phase cells, and cells following 12 hours of glucose withdrawal (Figure 5A). A previous study showed that trehalose levels dissipated immediately following dilution of stationary phase cells into growth media [31]. We similarly observed rapid kinetics of trehalose depletion in both spore and glucose-starved cells following their dilution into growth media (Figure 2A and 5B). In both cases, trehalose was essentially undetectable prior to cell division, arguing against its accumulation as accounting for the cellular memory of these states, which persist through multiple generations. Owing to its contributions to stress resistance independent of trehalose production, we genetically interrogated *TPS1* for a role in starvation induced cellular memory [29, 30]. Because *tps1∆* causes defects in growth in glucose containing media, we grew the cells for these experiments in YP+Galactose (YPG) and then subjected them to galactose withdrawal. We found that WT and *tps1∆* cells exhibited equivalent degrees of acquired stress resistance in response to galactose withdrawal Figure 5C). Interestingly, and as seen in a previous study, galactose starved *tps1∆* cells acquired the ability to make colonies on glucose containing YPD plates, though we were unable to detect any heritability of this capacity through liquid YPG cultures [28] (Figure 5D and data not shown). Collectively, our findings rule out a role for trehalose or of Tps1 in the cellular memory of spores or glucose starved cells.

**Figure 5.**
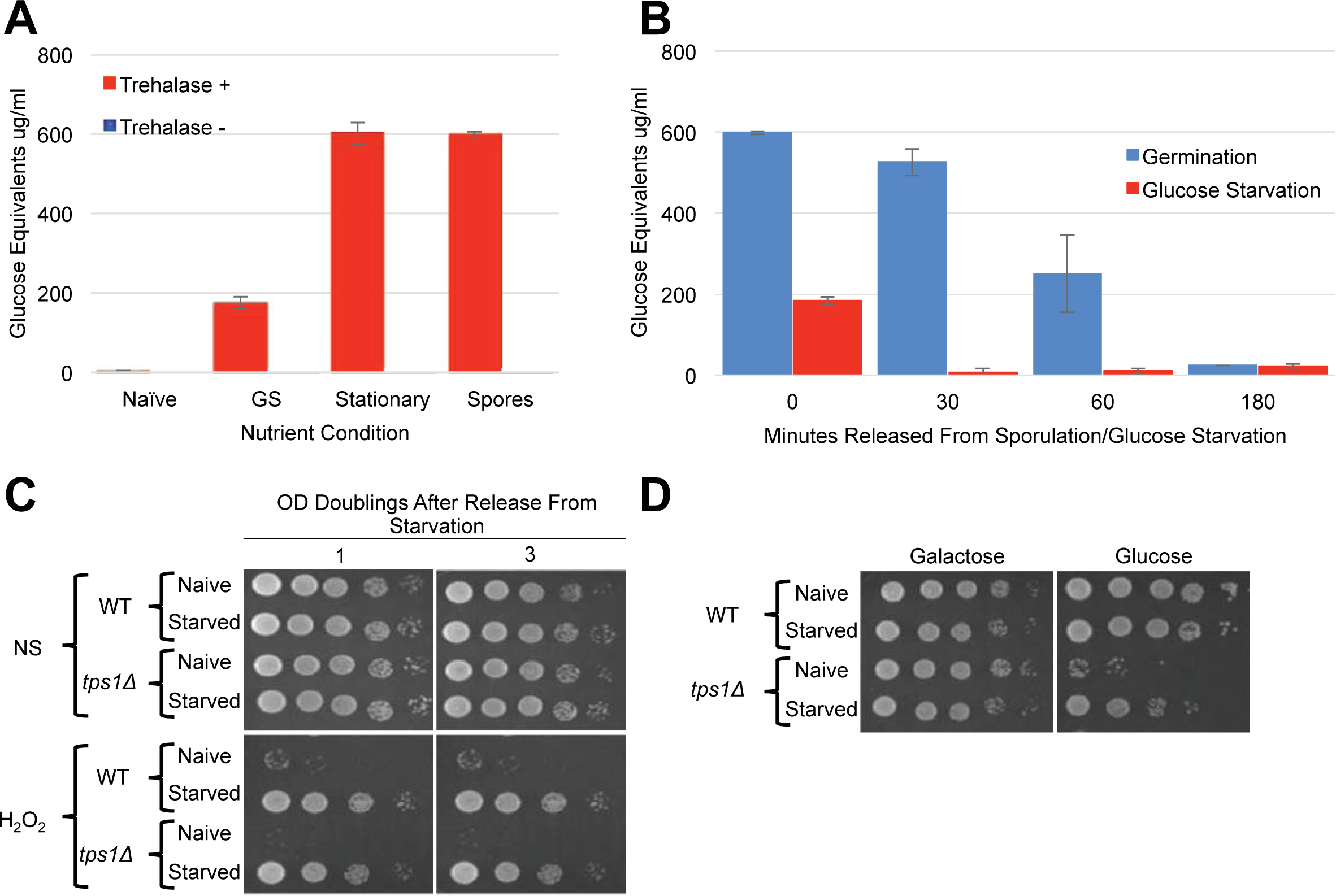
Neither trehalose accumulation nor *TPS1* activity accounted for cellular memory. A) The indicated cells were extracted and assayed for trehalose levels using an enzymatic assay. Error bars represent 1 standard error of the mean from 3 biological replicates. B) These same cells were diluted into YPD media and trehalose levels of the resulting cultured cells were determined at the indicated timepoints. Error bars represent 1 standard error of the mean from 3 biological replicates C) H_2_O_2_ stress resistance spot assay of 12 hour galactose starvation (Starved) and mid-log cells (Naive) of WT and *tps1Δ* are shown. NS represents cells that were not stressed with peroxide.D) 10-fold serial dilution spot assay of WT and *tps1Δ* on either SC-Galactose plates (Galactose) or SC-Glucose(Glucose) plates.

### Starvation induced cellular memory occurs independently of Msn2/4 activated programs

As mentioned above, the Msn2/4 transcription factors are essential for acquired stress resistance in response to varied inducing cues, and ectopic induction of an Msn2/4 target gene, *CTT1*, was sufficient to provide H_2_O_2_ resistance [9, 10, 32]. To compare starvation induced cellular memory with these other forms, we first evaluated *CTT1* induction using a strain that expressed Ctt1-GFP from its endogenous locus. Similarly to as reported previously, we observed robust *CTT1* induction in response to NaCl treatment, and this induction was prevented by thiolutin [10] (Figure 6A). In contrast, glucose starvation did not lead to any detectable Ctt1-GFP expression (Figure 6A). Curiously, we found that NaCl treatment of these glucose starved cells led to *CTT1* induction, which itself was also inhibited by thiolutin (Figure 6A).

**Figure 6.**
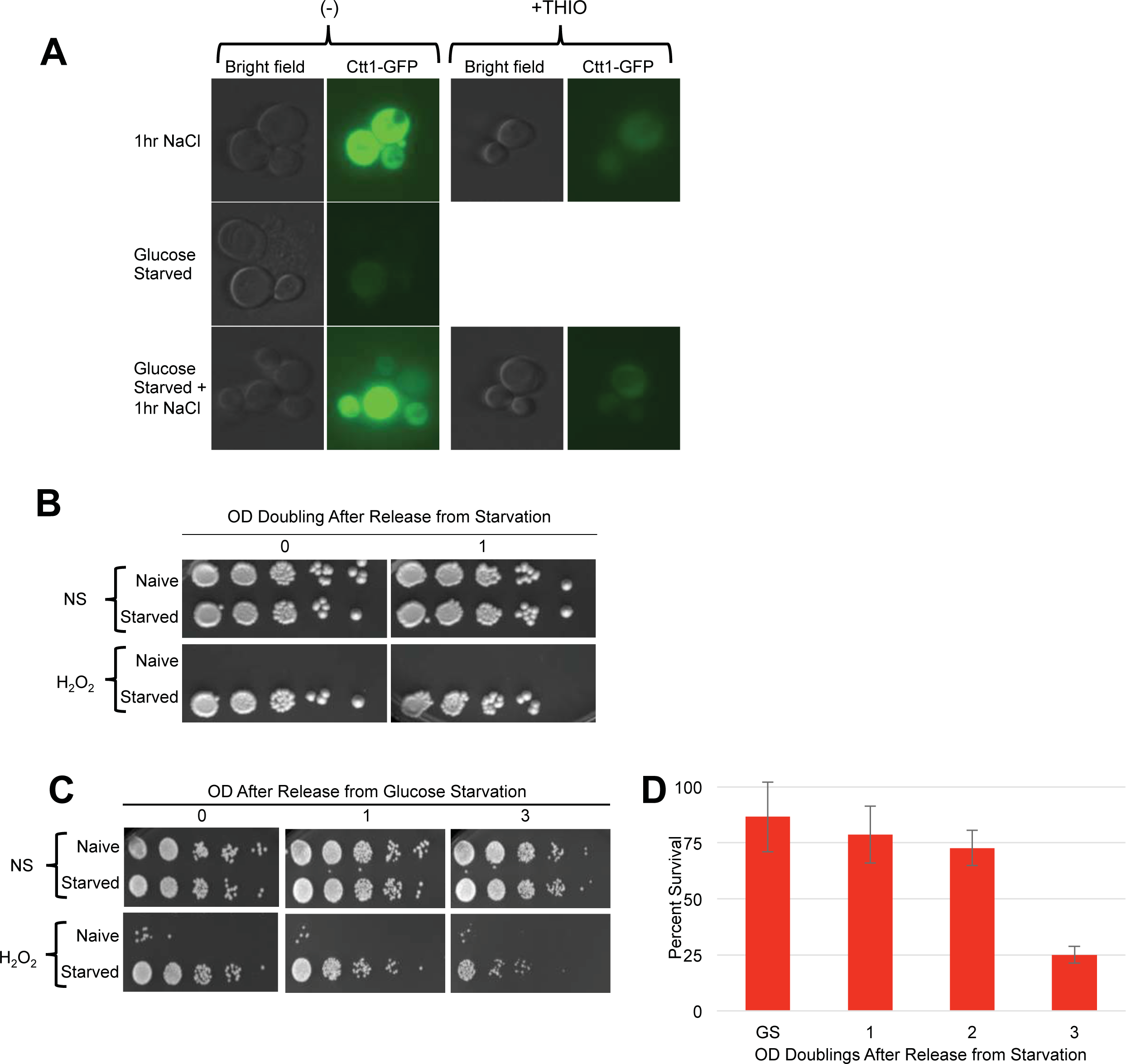
Ctt1 and Msn2/4 are not involved in starvation induced cellular memory. A) Representative images of a CTT1-GFP strain after 1 hour of NaCl exposure, 12 hours of glucose starvation (Starved), or 12 hours of glucose starvation + 1 hour of NaCl treatment (Starved + 1hr NaCl) are shown in the left panels. The right panels show the same cells treated with thiolutin concurrent with NaCl exposure. B) A *ctt1∆* strain was compared with an isogenic WT strain for starvation induced cellular memory. Survival of WT and *ctt1∆* cells to H_2_O_2_ exposure using spot tests is shown. C) H_2_O_2_ stress resistance spot assay of 12 hour glucose starvation (Starved) and mid-log cells (Naive) of a WT and *msn2Δ msn4Δ* strain. D) Survival of *msn2∆ msn4∆* cells released from glucose starvation and exposed to H_2_O_2_ was quantified. Error bars represent 1 standard error of the mean.

Our above finding was somewhat surprising, as glucose starvation is known to cause widespread transcriptional and translational dormancy [33, 34]. In this quiescent glucose starvation induced state, many mRNAs are thought to re-localize to P-body RNA storage organelles for utilization upon re-introduction of glucose, though the generality of this has been called into question [35, 36]. Our findings show that the glucose-starved cells mounted a gene expression response to NaCl that somehow over-rid their dormancy. Given thiolutin’s capacity for interfering with translation as well as transcription, it is difficult to determine if both of these steps are involved in *CTT1* induction in the glucose starved cells. At a minimum, it seems likely that *CTT1* mRNA that is stored in P-bodies in these glucose starved cells may be released and made available for translation by NaCl treatment, similarly to what has been suggested to occur in stationary phase cells [37]. Whatever the case may be, the absence of detectable Ctt1 in glucose starved cells is incompatible with a model in which its induction accounts for their cellular memory. To further substantiate this conclusion, we determined that deletion of *CTT1* had no obvious effects on starvation induced cellular memory (Figure 6B).

Msn2/4 activates many genes besides *CTT1* that might collectively contribute to starvation induced cellular memory [32]. Indeed, the nuclear localization of Msn2, and its presumed associated activation, is known to occur in response to glucose withdrawal, underscoring it as a candidate factor contributing to starvation induced cellular memory [38]. To more comprehensively interrogate the role of Msn2/4, we constructed an *msn2∆ msn4∆* double mutant strain for phenotypic analysis. We found that the deletion of *MSN2/4* had no consequence for glucose withdrawal mediated acquired stress resistance or cellular memory of this state (Figure 6C). We quantified the heritability of cellular memory in *msn2∆ msn4∆* mutants and found it to be indistinguishable from WT cells (Figure 6D).

## Conclusions

We describe here new examples of cellular memory in yeast that are exhibited in meiotically produced spores as well as glucose starved haploid cells. These cells exhibit resistance to multiple environmental stresses, and this heightened stress resistant state is inherited by newly produced daughter cells through at least one mitotic division. Moreover, the persistence of stress resistant cells over many ensuing divisions of these cultures suggests that these original cells retain this capacity for a very long time. Though we rule out roles for trehalose, *TPS1*, or of *MSN2/4* in them, our interrogation of the properties of these stress resistant states is nascent; further studies will be required to understand them. A plausible interpretation of our findings is that nutrient starvation underlies both the spore and haploid starved states. Indeed, in addition to glucose starvation, sustained starvation of haploid cells for nitrogen and phosphate also induce acquired stress resistance, though the cellular memory of these have not been investigated [16]. Further studies are likewise needed to distinguish these starvation-induced states and to assess their combinatorial consequences and relation to the cellular memory of meiotically produced spores.

The occurrence of cellular memory is widespread, seen in virtually all kingdoms of life, suggesting fundamental ancient origins. Stemming from this premise, it seems unlikely to us that gene expression programs and the derived epigenetic mechanisms regulating them can fundamentally account for such a phenomenon. As illuminated in the NaCl cellular memory studies interrogating Ctt1 [10], we suspect altered perdurance and inheritance of proteins likely underlies the cellular memories we describe here. If such a phenomenon indeed underlies these cellular memories, they must involve proteins other than Ctt1, necessitating the extension of whatever mechanisms led to Ctt1 enhanced perdurance and heritability to other proteins. An attractive speculation is that protein aggregation and/or some other widely acting modulation of the proteome underlies these cellular memories. Indeed, aggregation of the Whi3 protein underlies cellular memory of prolonged mating pheromone exposure in yeast, while induction of intrinsically disordered proteins explains the profound stress resistance exhibited by tardigrades [39, 40]. Whatever mechanism underlie the varied forms of starvation induced cellular memory we describe, their remarkable persistence seem well suited for screening methods to identify gene deletions impacting them using genomic approaches.

## Materials and Methods

### Yeast husbandry and strain construction

Strains were constructed using standard methods and listed in Table S1. Sporulation and germination studies were performed using a wild strain isolated from Oak tree exudates that sporulates to near 100% efficiency [41–43]. All other experiments utilized strains derived from the BY4742 background. Gene deletion strains were selected from the knockout collection and put through genetic crosses and tetrad dissections to engineer the strains used here [44]. The CTT1-GFP strain originates from the published GFP collection [45]. Sporulation and YPA growth were performed as described previously [42]. For germination studies, spores were then diluted at 0.15 OD_600_ in pre-warmed YPD at 30°C and incubated with shaking at 200 RPM. Germinating cells were not grown higher than 0.45 OD_600_ and when they reached these OD_600_ concentrations were diluted back to 0.15 OD_600_. O.D._600_ measurements were conducted every hour to track for doublings. Morphological changes were observed throughout germination using a microscope.

### Ether Sensitivity Assay

Ether sensitivity was tested by exposing spores and germinating cells to 33% ether and plating for survival as described previously [21]. Briefly, aliquots of culture containing sporulated cells or germinating/post-germination cells were removed from the culture and treated with 33% di-ethyl ether. At 2 minute intervals between 2-10 minutes of ether treatment, aliquots were removed,10-fold serially diluted in H_2_O, and 5 µl of these cell suspensions were spotted onto YPD plates. An assay was conducted every hour for the first 6 hours of germination timecourses. For drug treatments, germinating cultures were treated with 10 µg/ml thiolutin (T3450 SIGMA), 2 µg/ml cycloheximide (C7698 SIGMA), or DMSO controls respectively for the entire 6 hours.

### Stress assays

Strains were patched onto YPD plates from −80°C freezer stock and grown overnight. Cells were struck for single colonies on YPD plates and grown in YPD media as above. Overnight cultures were then diluted in fresh YPD media at an O.D._600_ 0.002. Cultures were grown for ~8 O.D. doublings to an O.D._600_ 0.3 at 30° C and then split into two separate cultures: one to be maintained in the “naïve” state, and one to be the recipient of an environmental perturbation. In order to maintain cultures in a naïve state, cells were diluted into fresh pre-warmed YPD at 0.15 O.D._600_ and were not allowed to grow beyond 0.45 OD_600_ before diluting back again. Unless specifically noted, in all acquired stress resistance and memory experiments, cells maintained as described above represent the naïve state. In a paired culture, cells were treated with various environmental conditions termed “treated cells”. To prepare cells for treatment, they were first harvested by centrifugation at 3000 RPM for 2 minutes and then washed with ddH_2_O twice with intervening harvesting by centrifugation. After wash with ddH_2_O, treated cells were harvested and then resuspended in pre-warmed YPD with NaCl added at a final concentration of 0.7 M stress for 1 hour or grown in YP media without a carbon source at 0.3 OD_600_ at 30°C. For sporulation experiments, cells were inoculated at 0.3 OD_600_ in YPA and for 12-14 hours to approximately 1.5 OD_600_ at 30°C. Terminally sporulated cells were produced by transfer of these “pre-SPO” cultures into sporulation media at an OD_600_ of 2, followed by incubation at 30°C for 24 hours with shaking at 200 RPMs. Carbon starvation was tested at 1 hour, 6 hours, and 12 hours. Treated cells were then harvested and washed in pre-warmed fresh YPD media. Spores were germinated in YPD media at 0.15 OD_600_ and not allowed to grow past 0.45 OD_600._ Germinating cells were maintained below 0.45 OD_600_ by serial dilutions to 0.15 OD_600._ Germinating cells were harvested and exposed to H_2_O_2_ ranging from 0-8 mM concentrations following 1, 2, 4, or 8 OD doublings of growth in YPD. Carbon starved cells were similarly released from starvation in stress-free YPD media for the indicated number of OD doublings. Equal numbers of cells, based on OD_600_ measurements, were harvested from naïve and treated cell cultures and then treated with 4mM hydrogen peroxide, 20% EtOH, or 3M NaCl for 2 hours in a 96-well plate. Cells were then 10-fold serial diluted and 5 µl of cells were plated on synthetic complete glucose plates or galactose plates. Afterwards, cells were grown for 48 hours at 30°C.

### RNA analysis

RNA was prepared using the hot acid phenol method. Total extractable RNA was then quantified using a spectrophotometer and normalized to the number of cells that produced it as approximated by input ODs of cells/spores.

### Trehalose Extraction and Measurement

10 O.D. pellets were collected and quickly washed with 1ml of ice cold H_2_O. Cell pellets were re-suspended in 0.25ml 0.25 M Na_2_CO_3_ and stored at −80°C until further processing. These cells were then boiled at 94-98°C for 4 hours, following which, 0.15 ml 1M acetic acid and 0.6ml 0.2 sodium acetate were added to each sample. Trehalose assays on these extracts were performed using a standard enzymatic assay as described previously [31].

## Acknowledgments

This work was supported by CIHR grant MOP-89996 to M.D.M. We are grateful to Dr. Charlie Boone and Dr. Brenda Andrews for providing the CTT1-GFP strain and to Amy Caudy for critical reading of the manuscript. The authors have no conflicts if interest.

---

**Figure S1.**
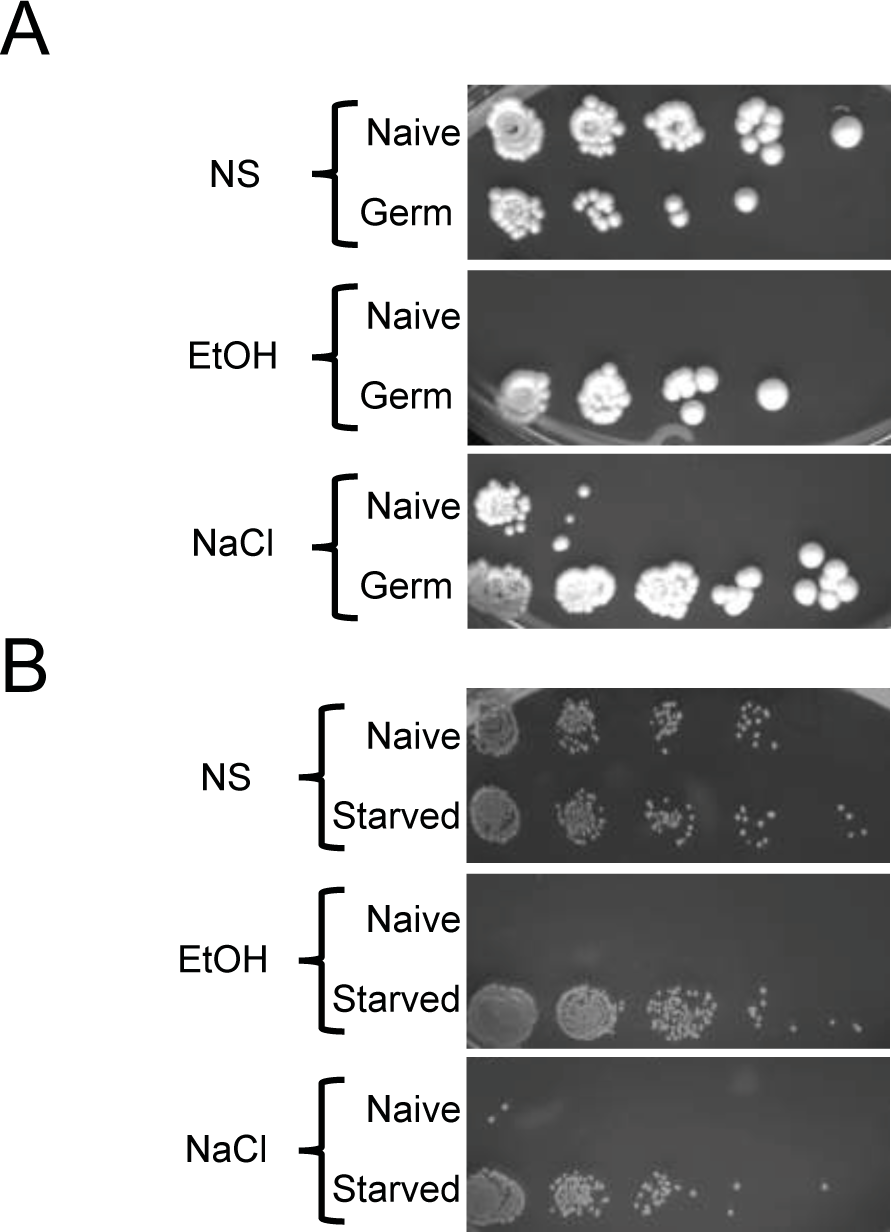
Germination and glucose starvation induce cross-stress resistance to ethanol and NaCl. A) 10-fold serially diluted spot assay of germinating cells treated with ethanol (EtOH) or NaCl. B) 10-fold serially diluted spot assay of 12 hour glucose starved cells treated with EtOH or NaCl.

**Table S1.**
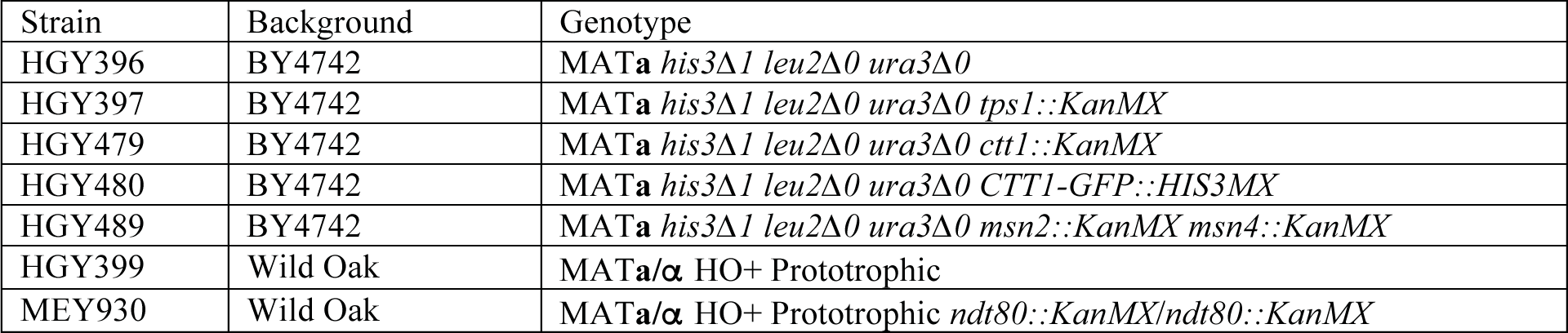

